# Predicting immune checkpoint therapy response in three independent metastatic melanoma cohorts

**DOI:** 10.1101/2024.05.01.592032

**Authors:** Leticia Szadai, Aron Bartha, Indira Pla Parada, Alexandra Lakatos, Dorottya Pál, Anna Sára Lengyel, Natália Pinto de Almeida, Ágnes Judit Jánosi, Fábio Nogueira, Beata Szeitz, Viktória Doma, Nicole Woldmar, Jéssica Guedes, Zsuzsanna Ujfaludi, Zoltán Gábor Pahi, Tibor Pankotai, Yonghyo Kim, Balázs Győrffy, Bo Baldetorp, Charlotte Welinder, A. Marcell Szasz, Lazaro Betancourt, Jeovanis Gil, Roger Appelqvist, Ho-Jeong Kwon, Sarolta Kárpáti, Magdalena Kuras, Jimmy Rodriguez Murillo, István Balázs Németh, Johan Malm, David Fenyö, Krzysztof Pawłowski, Peter Horvatovich, Elisabet Wieslander, Lajos V. Kemény, Gilberto Domont, György MarkoVarga, Aniel Sanchez

**Affiliations:** Department of Dermatology and Allergology, University of Szeged, Szeged, Hungary; Semmelweis University Department of Bioinformatics, Budapest, Hungary; Department of Pediatrics, Semmelweis University, H-1094 Budapest, Hungary; Section for Clinical Chemistry, Department of Translational Medicine, Lund University, Skåne University Hospital Malmö, 205 02 Malmö, Sweden; HCEMM-SU Translational Dermatology Research Group, Tűzoltó street 37-47. 1094 Budapest, Hungary; Department of Physiology, Semmelweis University, Tűzoltó street 37-47. 1094 Budapest, Hungary; Department of Dermatology, Venerology and Dermatooncology, Semmelweis University, Mária street 41. 1085 Budapest, Hungary; Clinical Protein Science & Imaging, Biomedical Centre, Department of Biomedical Engineering, Lund University, BMC D13, 221 84 Lund, Sweden; Chemistry Institute Federal, University of Rio de Janeiro, Rio de Janiero, Brazil; Department of Internal Medicine and Oncology, Semmelweis University, Budapest, Hungary; Department of Pathology, Albert Szent-Györgyi Medical School, University of Szeged, Állomás utca 1, Szeged H-6725, Hungary; Competence Centre of the Life Sciences Cluster of the Centre of Excellence for Interdisciplinary Research, Development and Innovation, University of Szeged, Dugonics tér 13, Szeged H-6720, Hungary; Hungarian Centre of Excellence for Molecular Medicine (HCEMM), Genome Integrity and DNA Repair Core Group, University of Szeged, Budapesti út 9, Szeged H-6728, Hungary; Therapeutics & Biotechnology Division, Korea Research Institute of Chemical Technology (KRICT), Daejeon, Republic of Korea; Research Centre for Natural Sciences, Institute of Molecular Life Sciences, Budapest, Hungary; Division of Oncology, Department of Clinical Sciences Lund, Lund University, Lund, Sweden; Chemical Genomics Global Research Lab, Department of Biotechnology, College of Life Science and Biotechnology, Yonsei University, Seoul, Republic of Korea; Department of Biochemistry and Biophysics, Karolinska Institute, Stockholm, Sweden; Section for Clinical Chemistry, Department of Translational Medicine, Skåne University Hospital Malmö, Lund University, Malmö, Sweden; Department of Biochemistry and Molecular Pharmacology, NYU Grossman School of Medicine; Department of Molecular Biology, University of Texas Southwestern Medical Center, Dallas, Texas, USA; University of Groningen, Analytical Biochemistry, Department of Pharmacy, Antinous Deusinglaan 1, 9713AV, Groningen, The Netherlands; First Department of Surgery, Tokyo Medical University, 6-7-1 Nishishinjiku, Shinjiku-ku, Tokyo 160-8402, Japan

**Keywords:** metastatic melanoma, immunotherapy, immunotherapy response, responders, non-responders, proteomics

## Abstract

While Immune checkpoint inhibition (ICI) therapy shows significant efficacy in metastatic melanoma, only about 50% respond, lacking reliable predictive methods. We introduce a panel of six proteins aimed at predicting response to ICI therapy. Evaluating previously reported proteins in two untreated melanoma cohorts, we used a published predictive model (EaSIeR score) to identify potential proteins distinguishing responders and non-responders. Six proteins initially identified in the ICI cohort correlated with predicted response in the untreated cohort. Additionally, three proteins correlated with patient survival, both at the protein, and at the transcript levels, in an independent immunotherapy treated cohort. Our study identifies predictive biomarkers across three melanoma cohorts, suggesting their use in therapeutic decision-making.

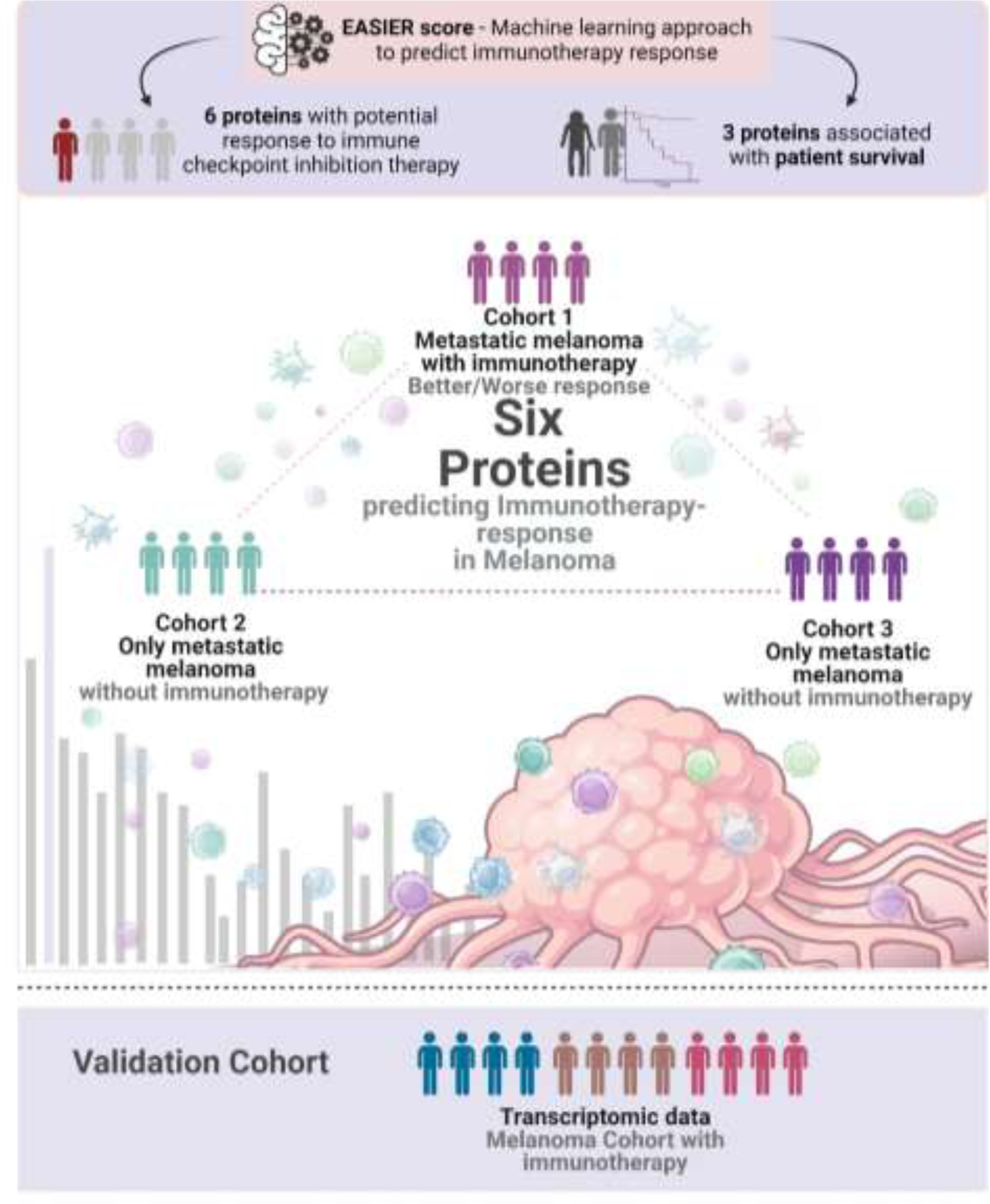

## Introduction

Cutaneous malignant melanoma is among the most therapy-resistant cancers with a high metastatic potential to distant organs.

Over the past decade, the treatment landscape of advanced and unresectable melanomas has been profoundly transformed, largely driven by advancements in our understanding of cancer biology and pathogenesis ^1–6^. This surge in knowledge has paved the way for innovative biological therapies, most notably immune-checkpoint inhibitors (ICI). However, the efficacy of ICI remains limited to a specific subset of patients, and the current clinical landscape lacks reliable biomarkers to assess the suitability of ICI therapy for individual patients.

Attributes of the melanoma sample could enhance the prediction of the immunotherapy response, such as; proteins related to antigen presentation ^4–6^, tumor mutation burden ^7^, CD8 protein in T-cells ^8^, the presence of tumor-infiltrating lymph cells ^9,10^, the composition of the tumor microenvironment (TME) ^11,12^, tumor burden ^13,14^, expression of self-antigens ^15^.Therefore, a critical unmet need in melanoma management remains: the identification of robust biomarkers capable of distinguishing responders from non-responders to ICI therapy in the early stage of melanoma progression.

Recently, a computational method predicting the immunotherapy response (ITR) of cancer patients was developed by quantifying signatures of the TME and its association with 14 different transcriptome-based predictors of anticancer immune responses. These predictors model different hallmarks of response to immune-checkpoint inhibitors ^16^. Based on this information, the authors constructed a machine learning model designated as the Estimate Systems Immune Response (EaSIeR) score, aimed at discerning the potential ITR in patients. The underlying algorithms for this model rely on RNA-seq data of the antitumoral immune response of 7,550 patients treated with PD-1/PD-L1 inhibitors across a spectrum of 18 solid tumors, including melanoma ^16^. Throughout this manuscript, we refer to the categorization that the EaSIeR score provides (ITR for responders and non-responders).

Here, we utilized data generated from the Human Melanoma Proteome Atlas project ^17,18^. Within the scope of this study, quantitative proteomics and comprehensive histopathological characterizations were conducted on 505 tumor samples, encompassing primary tumors and metastases across 26 organs. Building on this foundational work, we identified 401 potential biomarkers associated with ICI response ^12^ in the first cohort (Cohort 1), where immunotherapy data was available for 22 melanoma patients. In the present study, we assess the immunotherapy response association of these proteins in two independent cohorts of metastatic melanoma patients who have not received immunotherapy (Cohort 2 and Cohort 3). Our analysis involved predicting the patient’s potential immunotherapy response utilizing the EaSIeR scoring system. The overarching aim of our study is to elucidate the potential immune-response associations of these 401 protein candidates within two large, independent melanoma patient cohorts naive to immunotherapy, and validate the top candidate hits through transcriptomic datasets. (For detailed data about the clinical information of Cohort 1, Cohort 2, and Cohort 3, see **Table S3A, Table S3B, Table S3C and Materials and Methods**.)

## Materials and Methods

### i. The melanoma patient cohorts

In this study, we included three independent cohorts of metastatic melanoma patients. The first dataset, also referred to as Cohort 1, served as our discovery cohort to investigate proteins that potentially could predict the response to immunotherapy. We used this cohort as the groundwork for our study. Cohort 1 consisted of twenty-four melanoma samples from twenty-two patients, all of which had not received any prior immunotherapy treatment at the time of sampling, ensuring that the assessment of protein expression profiles occurred prior to any therapeutic interventions. Based on the progression data from patients undergoing immune checkpoint inhibitor (ICI) treatment, we defined two distinct subgroups: one characterized by progression (progressed subgroup) and the other by non-progression (non-progressed subgroup) during immunotherapy. Subsequently, we identified proteins within these subgroups that predict either better outcomes (no progression during immunotherapy, resulting in long progression-free survival) or worse outcomes (progression during immunotherapy, leading to short progression-free survival) in response to immunotherapy. The quantitative proteomics analysis led to the identification of a set of proteins that were significantly correlated with short or long progression-free survival and therefore were considered potential predictors of better or worse ITR. (Multiple Cox regression *p-value < 0.05). The detailed clinicopathological data of the cohort is presented in the supplementary material. (**Table S3A**) ^12^

The proteins that were identified in Cohort 1 were also examined in two independent melanoma cohorts (Cohort 2 and Cohort 3). Cohort 2 included 142 metastatic melanoma samples and also served as a first selection cohort in our analysis. At the time of sample collection, the patients had not received any prior treatment, and we have limited information about the subsequent application of immunotherapy in Cohort 2 patients. From the samples of Cohort 2 information of histopathology, and both quantitative proteomics and transcriptomics analyses were available. By proteomics and transcriptomics, 12,695 proteins and 11,468 genes were quantified. The resulting data from proteomics and transcriptomics served as a basis for the adjustment of protein scoring in our study. In Cohort 2, we utilized the EaSIeR scoring system to predict the responders and non-responders to immunotherapy based on the mechanistics signatures (e.g., immune cell quantification, pathway activity, transcription factor activity, ligand-receptor pairs, cell-cell interactions ^16^) which determine the responsiveness of the samples at protein level. By scoring the responder and non-responder samples, we were able to identify the proteins in both groups and compare with the proteins from Cohort 1. The comprehensive clinicopathological information about the aforementioned cohort is involved in the supplementary material (**Table S3B**). A manuscript describing this cohort is submitted to biorxiv.org. ^19^

Cohort 3, also referred to as a second selection cohort, consisted of a total of 44 metastatic melanoma samples. These patients had not undergone any immunotherapy treatment at the time of sample collection. Although we have information about the use of immunotherapy in this cohort, there is no available data on the immunotherapy response. From Cohort 3 quantitative proteomics information was available, all together 9040 proteins were identified and quantified by LC-MS/MS in Cohort 3. Therefore, we were able to correlate these proteins with transcriptome-based mechanistic signatures (e.g., immune cell quantification, pathway activity, transcription factor activity, ligand-receptor pairs, cell-cell interactions ^16^) which contribute to anticancer immune responses. Based on the correlation, ITR score could be investigated in the samples. The detailed clinicopathological information about the aforementioned cohort is described in the supplementary material **(Table S3C**).

### ii. Selection of proteins from a previously published dataset, testing and calculation of the predicted immune response in additional melanoma patients

Based on Szadai et al. (Cohort 1, ^12^) proteins associated with response to immunotherapy from 22 patients were selected to analyze this association in two new different cohorts (Cohort 2, Cohort 3). Firstly, the ITR scoring system, based on RNA-seq data proposed by Óscar Lapuente-Santana et al ^16^, which estimates the likelihood of patients responding to immune checkpoint blockade therapy was used. To test whether this scoring can be applied also at the protein level we used the Cohort 2 (142 patients) to correlate the predicted EaSIeR scoring obtained from RNA-seq data and the predicted EaSIeR scoring obtained from proteomic data (both datasets collected from the same cohort). The correlation was evaluated using a Pearson correlation test, with *p-values < 0.05 considered statistically significant. (**Table S2A, Table S2B, Table S3**). After observing a significant correlation between EaSIeR scores derived from RNA and protein levels in Cohort 2, we calculated the EaSIeR score for each of the 44 patients in Cohort 3. This allowed us to assign an ITR (immune checkpoint blockade therapy response) scoring to each patient in Cohort 3. (**Table S3D**)

### iii. Correlation between proteins linked to ITR and the scoring of ITR in patients

To determine which of the potential immune-response-related proteins selected from Cohort 1 ^12^ could be found in Cohorts 2 and 3, we first correlated the EaSIeR scores assigned to the patients in each cohort with the expression levels of each protein. Pearson correlations with adjusted p-values < 0.05 were considered significant. Second, patients from Cohorts 2 and 3 were stratified based on EaSIeR score (i.e., predicted ITR) to quartiles (Q1-Q4). The best potential responder patients were grouped in Q4 and poorest potential responders were in Q1. Next, a Student’s T-test was used to identify differentially expressed proteins (DEP) between poorest responders (Q1) and the best responders (Q4). Proteins with adjusted p-values (using Benjamini-Hochberg (FDR) method) < 0.05 were considered differentially expressed. (**Table S2A**)

All analyses described in **Figure 1**, **Figure 2** and **Figure 3** were performed using RStudio version 4.2.1. The summary results of the analysis can be found in **Supplemental material 3.** The code used for the analysis of this section is available in **Supplemental material 3**.

**Figure 1.**
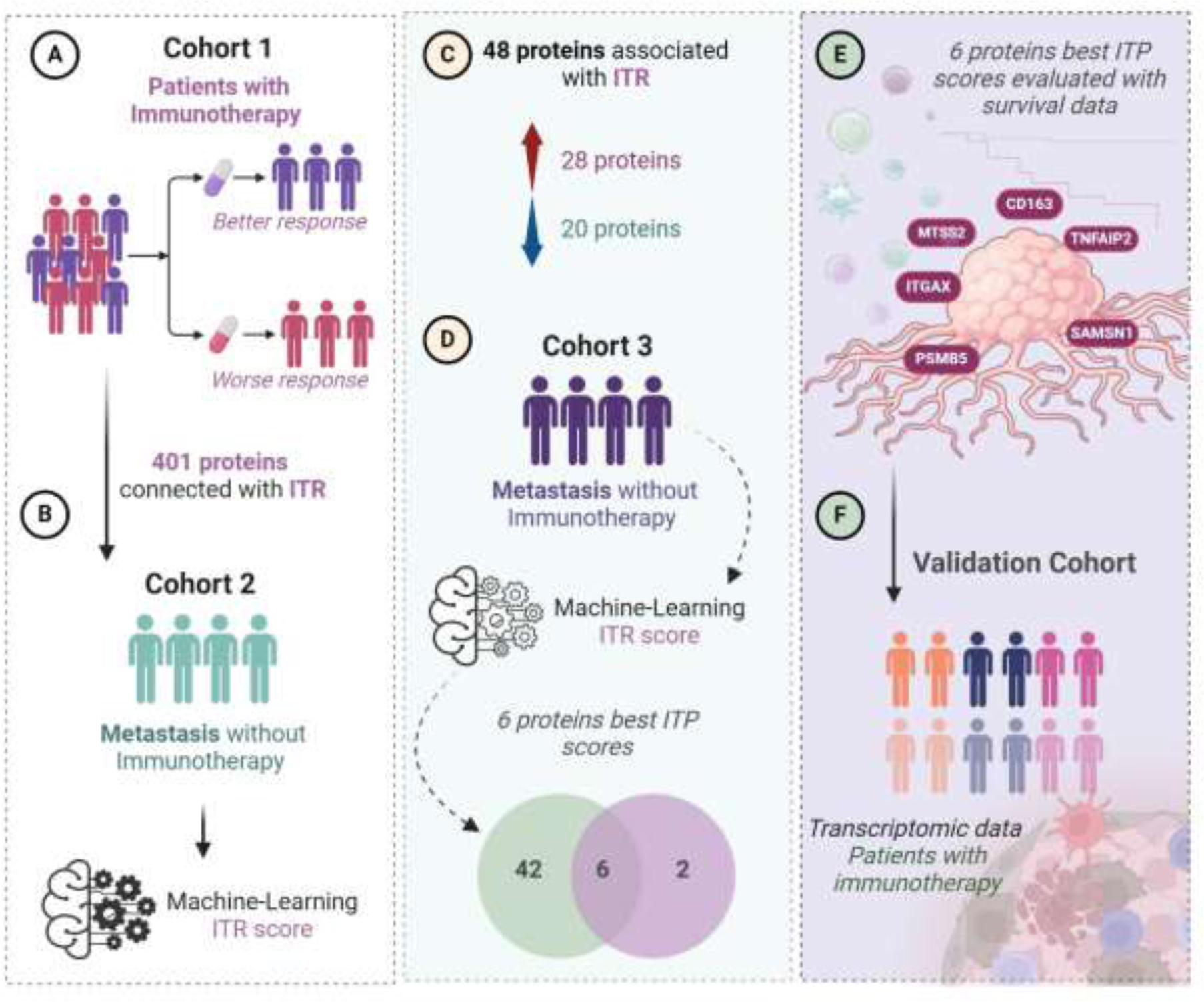
Workflow of the study step by step. From the first identified proteins through the testing of the ITR scoring until the final selection of the top six proteins. (A) 401 proteins were previously identified predicting ITR from a discovery cohort (Cohort 1, n=22). (B) Global proteomics outcomes were scored (ITR score) for immunotherapy response based on a machine-learning algorithm in a second independent cohort of patients with metastatic melanoma without immunotherapy (n=142). (C) 48 proteins were associated with ITR in Cohorts 1 and 2, 28 proteins upregulated and 20 downregulated. (D) Among these 48 proteins, 6 showed the best ITR score in Cohort 3, which is also composed of patients with metastatic melanoma (n=44). (E) correlation of the 6 proteins with the best ITR score with survival. (F) Validation using transcriptomics data in a cohort who received immunotherapy.

**Figure 2.**
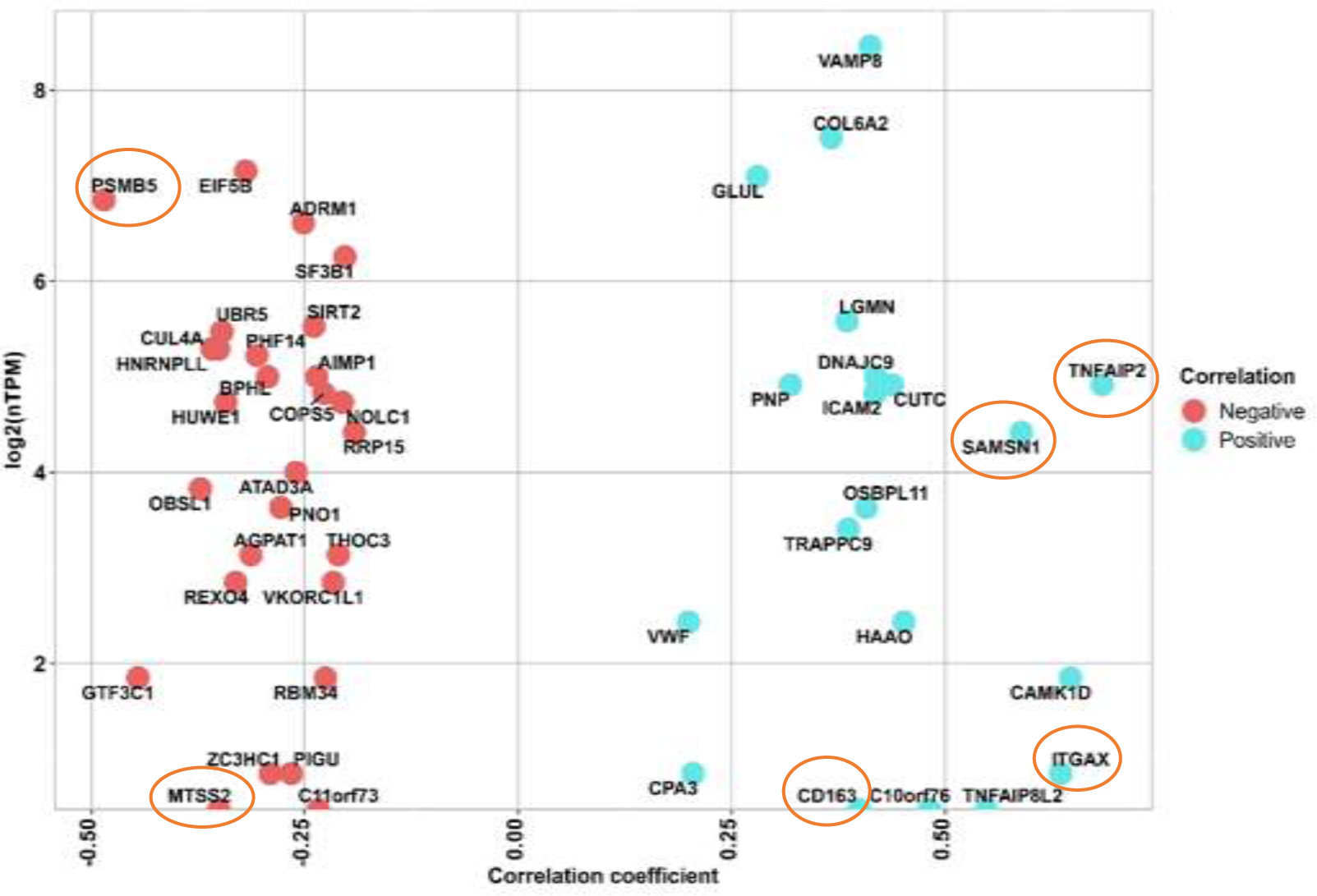
Scatter plot of the identified 48 proteins from Cohort 2. The y axis shows the RNA expression at single cell level (nTMP) values, x axis shows the correlation between protein intensity and ITR, and the identified 6 proteins marked with orange circle. Negatively correlated proteins are illustrated in red color, and positively correlated proteins are shown in blue color.

**Figure 3.**
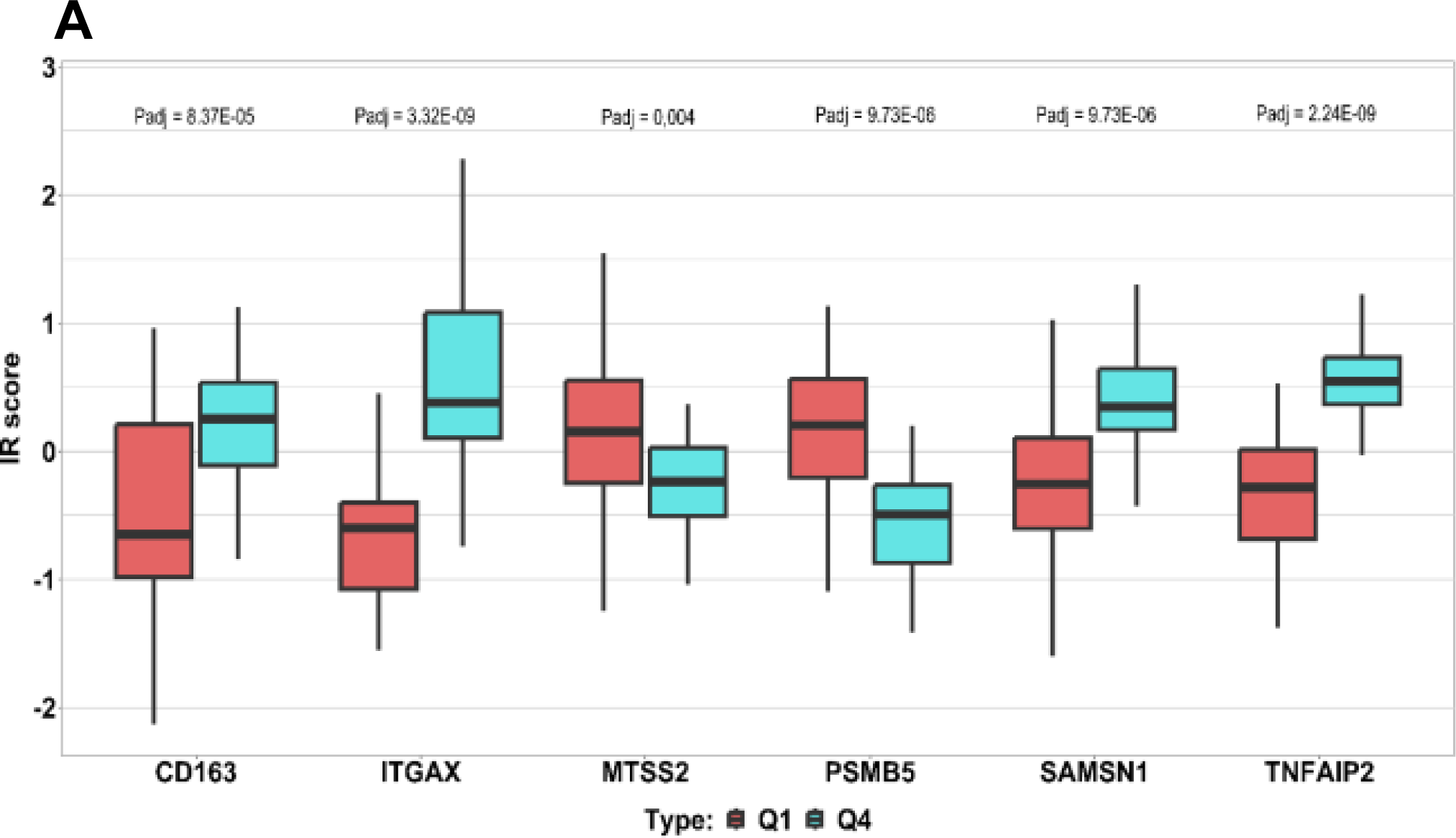
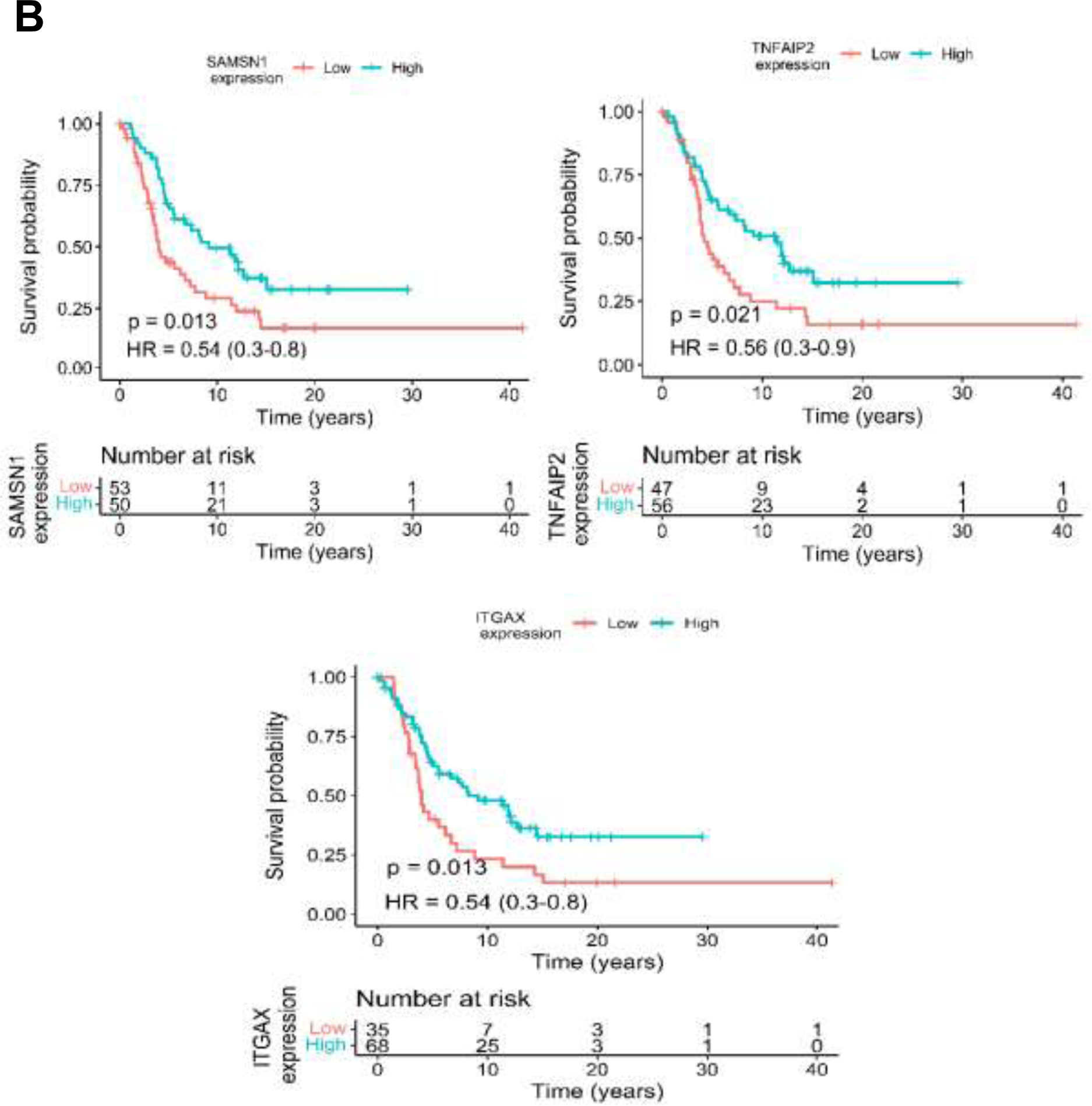
The differences of ITR scores in immunotherapy response and disparities in protein expression predicting survival. (A) represents ITR boxplot of the 6 proteins, Q1 vs Q4 values for each protein are shown to the left and the right side, respectively. Q1 representing potential non-responders is highlighted by a red box, while Q4 signifying potential responders are presented with a blue box. (*p-value < 0.05, Student-T test). (Supplementary Document S1) (B) Kaplan-Meier plots of the SAMSN1, TNFAIP2, and ITGAX, in patients where elevated expression of these proteins show a significant correlation with increased overall survival.

### iv. Selection of the proteins best associated with ITR and their Gene Ontology Enrichment Analysis

Proteins identified in Cohorts 2 and 3, which were found to be significantly associated with immune therapy response (ITR) (as observed by Szadai et al. ^12^) in **Figure 1**, were combined using Venn diagram and common proteins were considered for further analyses.

To investigate the molecular functionality of the proteins associated with ITR, we selected the list of proteins that were observed as significant in Cohort 2, as it was the largest list. We then conducted a Gene Ontology (GO) Enrichment Analysis (see **Table S2F and Table S3E**).

### v. Relationship of the top selected proteins with survival

Significant proteins found to be potentially related to ITR in both Cohorts 2 and 3 were considered the top significant proteins. To analyze the relationship of each of these proteins with survival, we used data from Cohort 2. First, a univariate analysis per protein was performed based on Kaplan–Meier (KM) curves. Secondly, we conducted Cox regression analysis to adjust the models for age at diagnosis, gender, tumor content of the sample, and clinical stage. To create Kaplan–Meier (KM) curves, proteins were categorized into low and high expression groups. This categorization was done by applying a receiver operating characteristic (ROC) curve per protein to detect the best cut-off point (based on Youden index) for discriminating between less or more than 2 years of survival. The protein expression (categorical variable) was used as the independent variable and the survival time of the patients served as the dependent variable. Patients whose protein values exceeded or fell below the cut-off point were categorized as having high or low protein expression, respectively.

### vi. Validation of the top proteins in transcriptomic cohorts with immunotherapy

Publicly available transcriptomic data obtained from melanoma tumors harvested before the initiation of PD1 inhibitor ^20^ and CTLA4 inhibitor ^14^ was used to investigate the top genes associated with progression-free and overall survival. Patients were ranked based on the gene expression of each gene. The survival of patients with the highest gene expression (top 10, 20 and 30% expression) was compared to survival of patients with the lowest expression) bottom 10, 20, 30%, respectively) for the Liu *et al.* dataset. To avoid selection bias, multiple cutoffs were used to avoid (see **Table S2E**). For the Van Allan dataset due to the lower number of samples, the top 25% and 50% were compared to the bottom 25% and 50%. The Log-rank test was used to determine significance.

### vii. Illustrations

The illustration showing the pipeline of the study (Figure 1) and the different molecular mechanisms involving the identified top proteins (Figure 5) were created with BioRender 2021 software ^21^. The Kaplan-Meier survival analysis was created by GraphpadPrism 8.0.1 (244)^22^.For the references Zotero Reference program was used. ^23^

## QUANTIFICATION AND STATISTICAL ANALYSIS

### **i.** Pearson correlations

For the correlation of the predicted EaSIeR scoring obtained from RNA-seq data and the predicted EaSIeR scoring obtained from proteomic data Pearson correlation test was used, with *p-values < 0.05 were considered statistically significant.

### **ii.** The comparison of Q1 and Q4

To compare the responders and non-responders in Cohort 2 and Cohort 3, patients were were stratified based on EaSIeR score (i.e., predicted ITR) to quartiles (Q1-Q4). Best responders were grouped in Q4 and the poorest responders were in Q1. Student’s T-test was used to identify differentially expressed proteins (DEP) between poorest responders (Q1) and best responders (Q4). Proteins with adjusted p-values (Benjamini-Hochberg (FDR) method) < 0.05 were considered differentially expressed.

### **iii.** Venn diagram

To filter the common proteins from Cohort 2 and 3, Venn diagram was utilized. (**Figure 1**)

### **iv.** Enrichment analysis

For the enrichment analysis for Gene Ontology Molecular Function (GOMF) plot from Cohort 2, we represented the top statistically significantly expressed proteins. Proteins with adjusted p-values < 0.05 were considered differentially expressed. This analysis was done using the R package clusterProfiler (version no. 4.4.4) ^24^, and ggplot2 (version no. 3.3.6) ^25^ for visualization. The code for the enrichment analysis can be found in **Table S3E**.

### **v.** Cox-regression survival analysis, Kaplan-Meier survival analysis and ROC curve

To investigate the relationship of each of these proteins with survival we used data from Cohort 2. The ROC curve was produced using IBM SPSS statistics package (26.0 version) software ^26^, with a significant threshold of P < 0.05 For the Kaplan-Meier survival analyses, we considered six proteins from the survival data of Cohort 2. These analysis were conducted for overall survival intervals (measured in years) and the calculations were based on models generated by the optimal cut-off value of each protein (**Table 3B, Table S2B, Table S2C, Table S2D, Figure S2, Figure 3B**). The KM curves were created using ‘ggsurvival’ and ‘ggsurvminer’ R packages. Additionally, Cox regression analysis was performed using ‘survival’ and ‘survminer’ R packages with p-values < 0.05 considered significant. Furthermore, the ROC analysis was conducted using SPSS 25 software (SPSS Inc, Chicago, IL, USA). (Table S2B, Table S2D and Supplemental material 3)

For the Kaplan-Meier survival analyses of the validation cohort we performed analysis for overall-survival, progression free-survival and disease-free survival, measured in years. (**Figure 4, Figure S3**) Patients in the Liu *et al.* dataset were categorized based on gene expression levels, with the top 10%, 20%, and 30% compared to the bottom 10%, 20%, and 30% for survival analysis. Multiple cutoffs were employed to mitigate selection bias (refer to **Table S2E**). In the Van Allan dataset, due to fewer samples, comparisons were made between the top 25% and 50% versus the bottom 25% and 50%. Significance was determined using the log-rank test.

**Figure 4.**
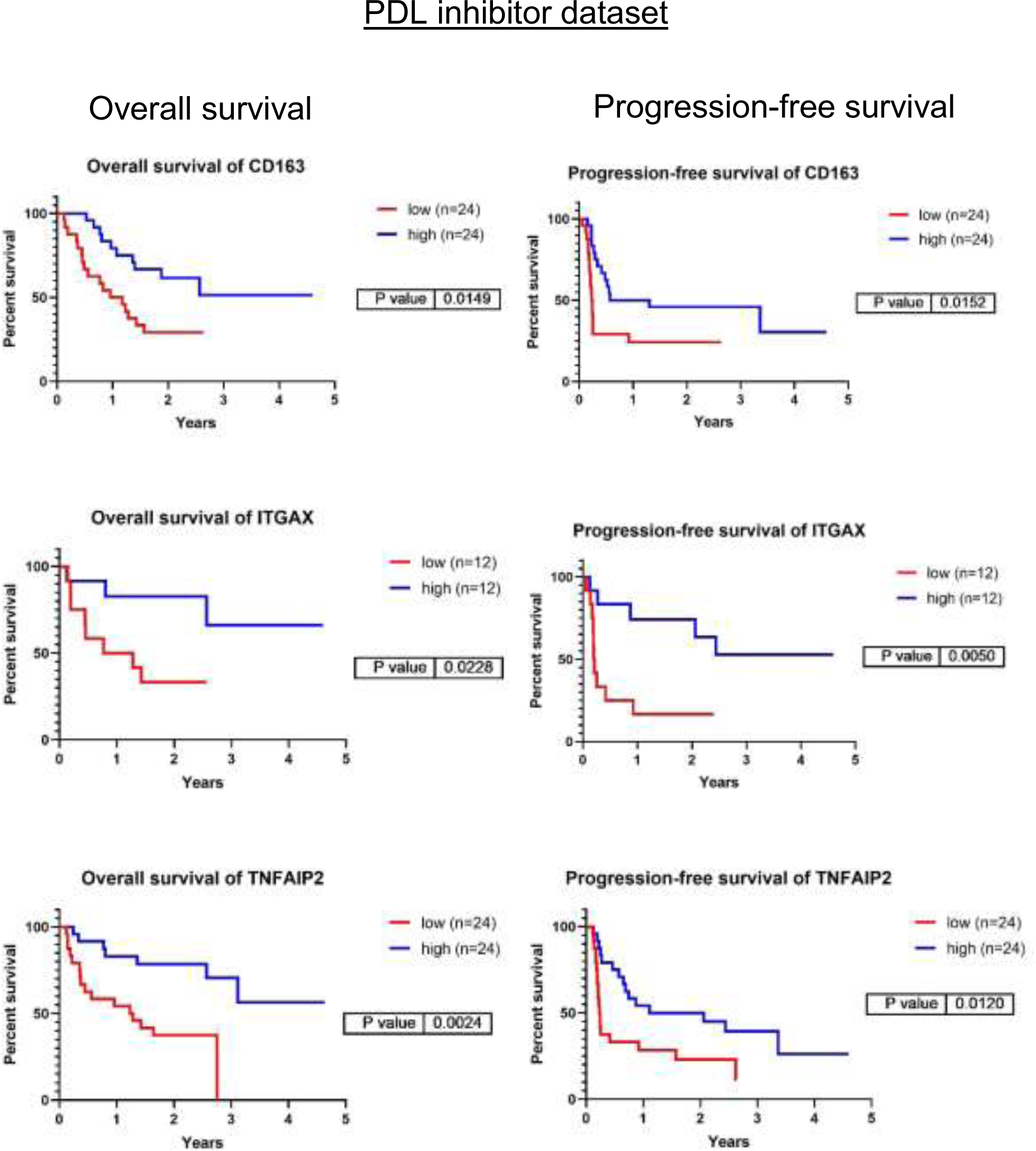
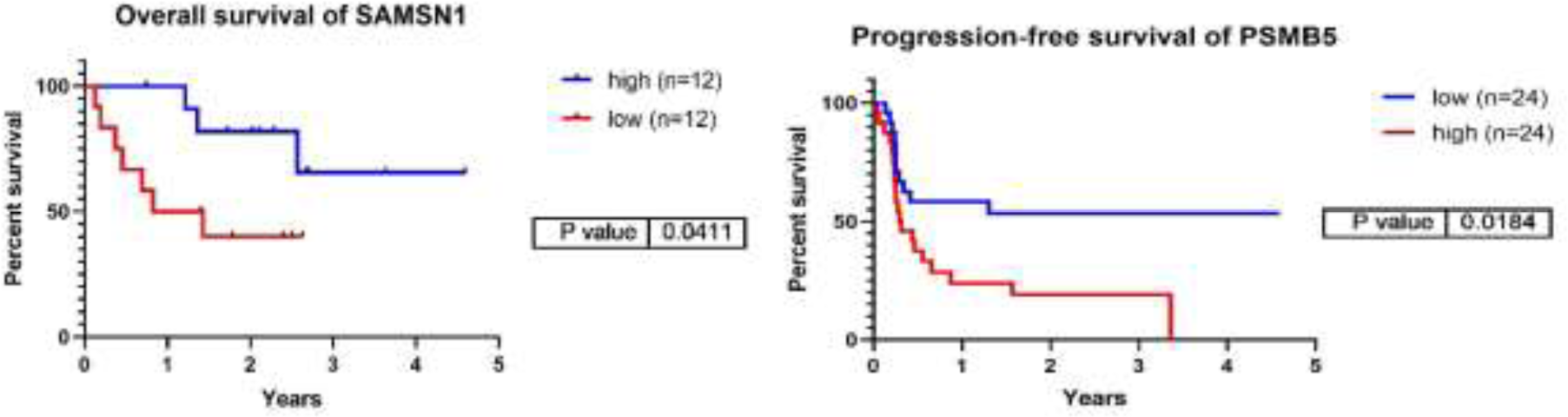
Kaplan-Meier survival analyses of the genes corresponding to the six top identified proteins focusing on overall-, and progression-free survival in the validation transcriptomic cohort of PD1 immunotherapy transcriptomic cohort. The cut off values used are the following: CD163 – PFS:20%, OS:20%, ITGAX – PFS10%, OS:10%, TNFAIP2 – PFS:20%, OS:20%, SAMSN1 – OS:10%, PSMB5 – PFS:20%. (PFS/Progression-free survival/,OS/Overall Survival/)

For the validation cohort, Kaplan-Meier survival analysis and figures including showing p-values, quartile values, mean values and 95% confidence intervals were produced was produced using Graphpad Prism 8. (**Figure 4, Figure S3**)

From the KM analysis, we extracted p-values based on Log-rank, Breslow, and Tarone-Ware tests. Proteins with a p-value < 0.05 in at least one of the three tests were considered significantly related to survival. (**Table S2B, TableS2C, Table S2D**)

### Materials availability

This study did not generate new unique reagents.

## Results

Selection of the immune checkpoint predictor proteins from Cohort 1 and the potential immune therapy response based on protein expression levels in Cohort 2 and 3

The three cohorts of metastatic malignant melanoma patients included in this study served multiple purposes: (a) the selection of potential immune-response-associated proteins (Cohort 1), (b) the identification and analysis of these proteins in treatment-naive cohorts (Cohort 2 and 3), (c) the exploration of their relationship with patient survival, and (d) their validation in independent cohorts treated with immunotherapy. The top candidates were further corroborated using transcriptomic data obtained from tumor samples in a cohort of patients that were administered either PD1 inhibitor ^20^ or CTLA4 inhibitor therapies ^14^. Tumor samples from Cohort 1-3 were included in the first human melanoma proteome atlas study ^17,18^. Detailed information concerning these cohorts and the data analysis workflow (**Figure 1**).

A total of 401 proteins were initially identified from Cohort 1 as potential immune-response-associated proteins. Within this metastatic cohort, 22 patients had undergone immunotherapy with varying degrees of treatment outcomes. Proteomic analyses conducted on samples from Cohort 1 ^12^ revealed that these 401 proteins were significantly correlated with progression-free survival, as evidenced by multiple Cox regression analyses (*p-value < 0.05), thereby positioning them as potential predictors of immunotherapy efficacy.

In order to expand the implications of these findings to Cohorts 2 and 3, we initially assess the utility of these proteins as potential indicators of immunotherapy response within these treatment-naive cohorts.

Considering that the samples from these cohorts were treatment-naïve at the time of evaluation, a non-conventional approach was employed to estimate the potential immunotherapy response in patients from Cohorts 2 and 3. Utilizing the EaSIeR scoring system (ITR scoring), we first estimated the ITR of 142 samples from 119 patients in Cohort 2. A statistically significant correlation (Pearson correlation test, *p-value < 0.0001, r = 0.7) (**Figure S1, Table S2A**) was observed between transcriptomics and proteomics data across these samples, enabling us to estimate the ITR of patients from Cohort 2 and 3 based proteomics data. After computing the estimated immunotherapy response score, based on protein expression profiles in samples from both Cohort 2 and 3, we further examined if the previously identified 401 proteins could be associated with the immunotherapy response scores in these independent, non-treated cohorts (Cohort 2 and 3).

### Significant proteins associated with ITR

After performing a Pearson correlation between the abundance profiles of each of the 401 previously identified proteins and the immunotherapy response scores in patients from Cohort 2, we found 48 proteins that exhibit significant correlation (*p-value < 0.05). Within this subset, 20 proteins were positively correlated with ITR (Pearson correlation coefficient (r), r (0.684) > 0), while 28 were negatively correlated (Pearson correlation coefficient (r), r (-0.485) < 0). A similar analysis performed on data from Cohort 3 resulted in eight proteins with significant correlation (*p-value < 0.05), two of them were negatively correlated with ITR, and six were positively correlated. To see the proteins that exhibited significant upregulation and downregulation, they were displayed at the single-cell level, based on the expression of RNA representing the production of these proteins in melanocytes. (**Figure 2, Table S1, Table S2A, Table S3D, Table S3E**).

Based on this analysis, we opted to prioritize six proteins (ITGAX, SAMSN1, CD163, TNFAIP2, MTSS2, PSMB5) as they emerged as the foremost candidates by virtue of their significant correlation to immunotherapy response in both Cohorts 2 and 3.

### Impact of the six proteins on ITR prediction

To determine the influence of the six identified proteins on ITR prediction within the two distinct cohorts (Cohort 2 and 3), we categorized the ITR into quartiles (Q1-Q4). The magnitude of the difference between the upper Q4 and lower Q1 quartiles served as a proxy for the protein’s predictive power in distinguishing the responsiveness to therapy. In our analysis, all six proteins demonstrated significant differences between Q1 and Q4 (Student T test, *p-value < 0.05) (depicted in **Figure 3 A, Table S2A**). Specifically, ITGAX, SAMSN1, TNFAIP2, and CD163 proteins were upregulated and associated with higher scores in potential responders. Conversely, MTSS2 and PSMB5 showed lower scores and were downregulated in the same group. As expected, these findings are in concordance with the trends observed in the previous Pearson correlation analysis (**Table S2A, Table S2B**).

### Association between the selected six proteins and survival Kaplan-Meier Univariate analysis

We analyzed the impact of these proteins on overall survival. The survival information was obtained from 127 patients of Cohort 2. The Kaplan-Meier curves unveiled that different levels of ITGAX, SAMSN1, and TNFAIP2 proteins were significantly associated with 2-year survival (depicted in **Figure 3 B**) (Cox regression, *p-value < 0.05). On the other hand, proteins CD163, PSMB5, and MTSS2 did not significantly associate with 2-year survival (**Figure S2, Table S2B, Table S2C, Table S2D**).

### Independent survival prognostic values of the selected six proteins

Furthermore, we delved into the prognostic survival values of these proteins in relation to other clinical parameters (e.g., age at diagnosis, gender, tumor content (%), and disease stage) within Cohort 2. Cox-regression models were created based on the clinical parameters (Cox model 1: associations with survival, Cox model 2: associations with Cox model 1 plus age at diagnosis, gender, and tumor content (%), Cox model 3: associations with Cox model 2 plus disease stage) and the independence of the identified proteins was analyzed from the clinical parameters. Interestingly, distinct outcomes emerged from the Cox regression survival analyses (Cox models 1, 2, and 3). ITGAX, SAMSN1, and TNFAIP2 were significant favorable predictors of survival from Cox model 1 (Cox regression, *p-value <0.05). Involving other clinical parameters such as age at diagnosis, gender, and tumor content (%) in Cox survival analysis model 2, ITGAX and SAMSN1 showed significant values. Three proteins (TNFAIP2, SAMSN1, CD163) for the third Cox survival analysis model represented significant independence from the clinical parameters, meaning that these proteins may be survival predictors regardless of clinical parameters such as gender, age at diagnosis, tumor content (%) or disease stage. Notably, when considering the investigated clinical parameters, SAMSN1 protein appeared as the most independent survival predictor considering the investigated clinical parameters (**Table 1, Table S2B, Table S2C, Table S2D)**.

**Table 1.** The identified proteins with their corresponding genes, accompanied by the outcomes of Cox regression survival analysis. Cox model 1 consists of survival, and Cox model 2 involves Cox model 1 along with age at diagnosis, gender, and tumor content (%). Cox model 3 includes all the parameters from Cox model 2 plus the disease stage. (Cox regression analysis *p-value < 0.05 considered as significant marked with *.)

| Protein | Gene | Cox model 1: cox<br>ph(surv(times,died)~gene) | Cox model 2: Cox<br>model 1+age<br>diag.+gender+ TC | Cox model 3: Cox model<br>2+Disease Stage |
| --- | --- | --- | --- | --- |
| P20702 | ITGAX | p < 0.05 (HR=-0.314)* | p < 0.05 (HR=-0.313)* | p = 0.084 (HR=-0.257) |
| Q03169 | TNFAIP2 | p < 0.05 (HR=-0.669)* | p = 0.059 (HR=-0.721) | p < 0.05 (HR=-0.637)* |
| P28074 | PSMB5 | p = 0.065 (HR=0.349) | p = 0.059 (HR=0.388) | p = 0.061 (HR=0.404) |
| Q9NSI8 | SAMSN1 | p < 0.05 (HR=-0.619)* | p < 0.05 (HR=-0.656)* | p < 0.05 (HR=-0.569)* |
| Q86VB7 | CD163 | p = 0.067 (HR=-0.303) | p = 0.059 (HR=-0.372) | p < 0.05 (HR=-0.414)* |
| Q765P7 | MTSS2 | p = 0.961 (HR=0.009) | p = 0.059 (HR=0.026) | p = 0.431 (HR=-0.155) |

### Validation of the identified proteins using transcriptomic data from patient cohorts undergoing immunotherapy

We selected the top 6 proteins that displayed a significant association with survival above and investigated whether the transcript levels of the genes encoding these proteins could predict the prognosis of patients receiving PD1 inhibitor ^20^ or CTLA4 inhibitor ^14^ therapies. Among the 6 genes investigated, 5 demonstrated a significant prognostic association (summary of all genes in all datasets tested displayed in **Table S2E**). Expression of CD163, ITGAX and TNFAIP2 were associated with better PFS and OS in response to PD1 inhibitor therapies (Log rank test, *p-value <0.05). ITGAX and TNFAIP2 were associated to both PD1 inhibitor and CTLA4 inhibitor therapies, whereas high expression of PSMB5 was linked to poorer PFS following PD1 inhibition (**Figure 4, Figure S3, Table S2E**). SAMSN1 gene exhibited a weak association with overall survival in the PD1 inhibitor therapy dataset, whereas MTSS1L did not demonstrate a connection with either progression-free or overall survival in the two datasets tested. These results collectively support our earlier findings that expression of CD163, ITGAX, TNFAIP2 and SAMSN1 may be linked to positive responses to immune-based therapies, whereas PSMB5 expression may correspond to resistance against such therapies.

### Biological function of the identified biomarkers

We investigated the molecular functions of the initial set of 48 proteins that displayed association with immunotherapy response in Cohort 2. Based on the Gene Ontology Molecular Function (GOMF) enrichment analysis, we observed that ITGAX as well as ICAM2, and VWF were linked to integrin binding. PSMB5 was found to be involved in the threonine-type endopeptidase activity, while TNFAIP2 and VAMP8 were implicated in SNARE binding, implying cell communication across membranes and exocytosis ^27^ (This is summarized in **Figure 5** and detailed in **Table S2F, Table S3E**). Furthermore, utilizing the mRNA expression values (nTPM) at single cell level as published in The Human Protein Atlas ^28^, we determined that ITGAX, TNFAIP2, SAMSN1, and PSMB5 exhibit high mRNA levels in melanocytes. Conversely, the mRNA expression of CD163 and MTSS2 in melanocytes is absent, implying that the source of these proteins could be from other cells in the microenvironment (for more detailed information, please refer to **Table S2F, Table S3E**).

**Figure 5.**
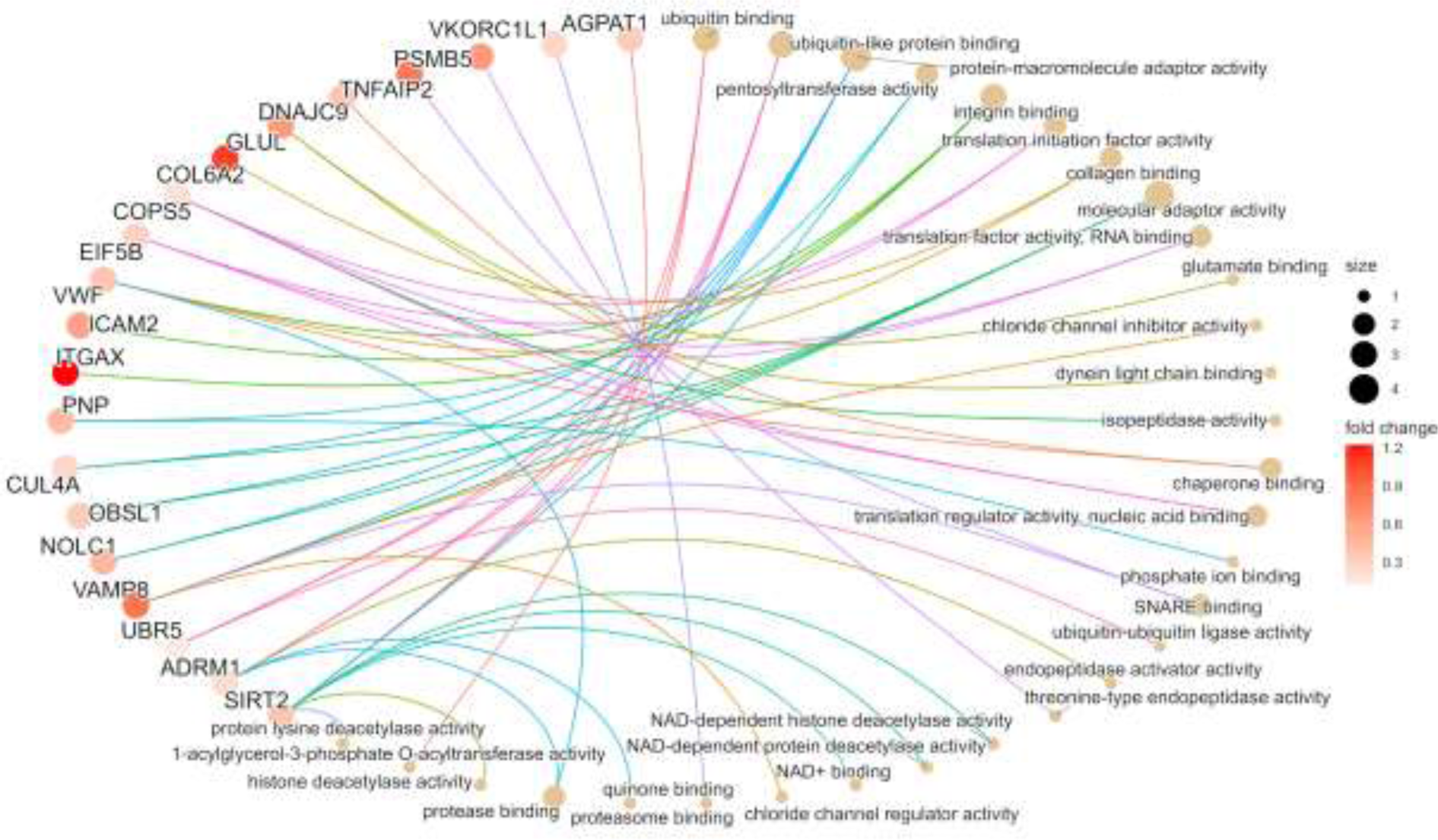
The scheme illustrates the Gene Ontology Molecular Function (GOMF) plot of the top 20 proteins from Cohort 2 based on the enrichment analysis. The image illustrates the most relevant molecular pathways connected to the first selected 48 proteins. In the case of pathways, the size of circles indicates the number of proteins involved in the functions. The proteins are presented in colors based on the magnitude of fold change.

## Discussion

Identifying suitable biomarkers to predict immunotherapy response presents a significant challenge within melanoma research and clinical practice. Currently, in the case of advanced melanoma patients, molecular-level assessment of the BRAF mutation status guides the selection of kinase inhibitors. However, the efficacy of targeted therapy remains uncertain owing to intricate mechanisms of resistance ^29^. Additionally, the expression of PD-L and PD- L1 proteins can exert an impact on the response within the tumor microenvironment (TME) enhancing the ability of immune cells to counteract undesirable signals originating from melanoma cells ^30^.

Although PD-L1, and PD-1-based immunotherapies currently serve as first-line treatments for advanced metastatic melanomas, the emergence of resistance remains a prevalent concern ^29^. Other approaches, such as digital imaging, have been explored for assessing PD-1/PD-L1 expression within melanoma tumor samples, aiding in identifying suitable therapy composition and timing^31^. Moreover, factors such as PD-1/PD-L1 expression, tumor stage, driver mutation status, and metastatic extent can offer valuable insights into the prognosis ^31^.

Due to the complexity of molecular pathways influencing melanoma progression, therapy response, and resistance, identifying a solitary protein biomarker that mirrors these events is challenging. In our study, we leveraged a previously published dataset, the Metastatic Melanoma Cohort Study ^17,18^, which offers proteomic and clinical insights from 263 primary and metastatic melanoma samples. Amidst the diverse patient cohorts in the Metastatic Melanoma Cohort Study ^17,18^, we narrowed our focus to individuals who received immunotherapy and exhibited a treatment response. To manage the extensive array of proteins within the immunotherapy patient pools, we employed a machine learning tool to predict the Immune-Therapy Response (ITR) ^16^. The scoring system incorporates all information on cell compartments from the melanoma samples and gives a score that may help in decision-making regarding therapy response. Through an extensive proteomic analysis conducted across three independent cohorts we identified six candidate proteins. Notably, the protein levels of ITGAX, SAMSN1, MTSS1L, PSMB5, TNFAIP2, and CD163 exhibited a correlation with the predicted ITR. Moreover, CD163, TNFAIP2, and SAMSN1 displayed a robust association with survival outcomes that remained significant independently of key clinical parameters such as gender, age at diagnosis, tumor content (%), and disease stage. Moreover, when delving into published transcriptomic databases containing immunotherapy-related information, we observed a substantial correlation between the expression pattern of the identified proteins and both overall and progression-free survival.

Furthermore, the six proteins that were identified exhibit association with distinct mechanisms. ITGAX (Uniprot: P20702), also known as CD11c serves as a receptor for fibrinogen and has an important role in cell adhesion mechanisms ^32^. This function likely extends to the tumor microenvironment (TME), where it participates in cell adhesion modulator ^33^, and cell-cell interactions during inflammatory responses ^34^ and is also produced in lower amounts by melanocytes ^35^. The ITGAX subunits are one of the most widely overexpressed proteins in various cancers ^36^, rendering them potential targets for antitumor therapies ^37^. Macrophages and T-cells are enriched with ITGAX. However, there are debated findings regarding integrin subunits in the context of melanoma, they may be related to the pathological stage, disease-free survival and melanoma metastasis. Moreover, a strong correlation has been established between the expression of integrin subunits and immune cell infiltration ^38^. These observations provide a plausible rationale for the positive role attributed to ITGAX. In our dataset, we observed an upregulation of ITGAX in parallel with better survival and enhanced therapy response. The transcriptomic data further strengthen this notion, as the ITGAX gene exhibited significant overexpression within the long-survival group marked by prolonged PFS and OS in both the PD1 inhibitor and CTLA4 inhibitor datasets. This consistency underscores the robust predictive potential of ITGAX. Further, treatments targeting integrin subunits have been successfully employed against various diseases. Notably, FDA-approved monoclonal antibodies, such as etrolizumab, have been deployed to obstruct CD11a units and ITGB7 (integrin subunit beta 7) in inflammatory diseases such as severe plaque psoriasis, Crohn’s disease or ulcerative colitis, respectively ^39,40^. Based on our results, we hypothesize that the role played by ITGAX holds promise in predicting the response to immunotherapy in melanoma.

The SAMSN1 (SAM domain, SH3 domain, and nuclear localization signals 1 ^41^ Uniprot: Q9NSI8) protein was also identified in patients exhibiting enhanced therapy response and higher survival rates. Interestingly, we observed a substantial upregulation in the expression of SAMSN1 gene related to overall survival in patients who received PD1 inhibitor therapy. SAMSN1 is implicated as a negative regulator in B-cell activation ^41^. Addition to this notion, Jönsson et al. underscored the association of the CD20-positive B-cell subset with a favorable prognosis for patients diagnosed with metastatic melanoma ^42^. In addition, Helmank *et al.* found that responders to neoadjuvant immunotherapy exhibited elevated levels of a predetermined B-cell signature in both baseline and early on-treatment samples ^43^. Moreover, in vitro studies have demonstrated that SAMSN1 contributes to the downregulation of cell proliferation and is also synthesized by melanocytes ^28,44^. As a tumor suppressor gene, the decreased expression of SAMSN1 was found in several cancers ^45^. Additionally, low SAMSN1 protein production in hepatocellular carcinoma and gastric cancer was associated with decreased overall survival and expanded tumor size ^45–47^. Thus, our results are in line with recent publications advocating that the levels of SAMSN1 protein can be associated with better immunotherapy response.

The TNFAIP2 (TNF alpha-induced protein 2, Uniprot: Q03169) was identified in our study as a predictor of better therapy response and was also upregulated in melanoma patients in correlation with long survival both in proteomic and in PD1 and CTLA4 immunotherapy transcriptomic cohorts. TNFAIP2 protein is known as an important player in inflammation, angiogenesis, proliferation, and migration and it is a cancer-related gene ^48,49^. The TNF alpha-induced protein 2 is produced mostly by lymphocytes, macrophages, mast cells in inflammation, and melanocytes ^48,50^. The TNFAIP2 expression can exhibit variation across different cancer types. For instance, in a recent publication, there was a comparison of normal and tumor tissue for mRNA variants of TNFAIP2 among all the cancers in the TCGA database. Lin Jia et al. discovered that TNFAIP2 mRNAs were upregulated in renal clear cell carcinoma, while in skin cutaneous melanoma, a contrasting pattern emerged where TNFAIP2 mRNAs were downregulated ^48^. Furthermore, a Kaplan-Meier survival analysis revealed that high TNFAP2 mRNA expression correlated with extended survival ^48^, aligning with the outcome observed in our study at the proteomic level.

The PSMB5 or proteasome 20S subunit beta 5 (Uniprot: P28074), demonstrated a correlation with decreased expression alongside improved survival and a positive response to immunotherapy. At the transcriptomic level, a significantly decreased gene expression was observed during progression-free survival in the PD1 immunotherapy dataset. The functional role of PSMB5 is intricate, primarily revolving around proteolytic functions, including degrading ubiquitinated proteins in the cell. Moreover, PSMB5 is produced by melanocytes, further accentuating its relevance in the context of melanoma ^51,52^. FDA-approved anti-proteasome agents like Bortezomib are used to treat multiple myeloma and mantle cell lymphoma in which the proteasome activity is high and associated with oncogenic functions ^53^. Wei et al. also demonstrated in triple-negative breast cancer that PSMB5 is an indicator of poor prognosis and the silencing of the PSMB5 gene can increase the sensitivity of breast cancer cells to chemotherapy and subsequently to apoptosis ^54^. Our results are in strong agreement with existing literature, suggesting that decreased PSMB5 protein expression might serve as a marker for enhanced prognosis in melanoma immunotherapy response.

Lastly, CD163 is an acute phase-regulated receptor involved in protecting tissues from free hemoglobin-mediated oxidative damage ^55^ (Uniprot: Q86VB7), and MTSS2 (MTSS I-BAR domain containing 2, Uniprot: Q765P7) is related to tumor metastasis and cancer progression via interactions with the actin cytoskeleton, and belongs to the MTSS family ^56^. Notably, both identified proteins seem not to be produced by melanocytes ^57,58^. Our findings showed an upregulation of CD163 associated with improved immunotherapy response and improved survival. Moreover, the gene expression of CD163 was significantly upregulated in the long-survival group (prolonged progression-free and overall survival) in the PD1 and CTLA4 immunotherapy transcriptomic cohorts. Contrary to our data, a recent publication presented that CD163+ tumor-associated macrophages in melanoma were positively correlated with deeper Breslow level, advanced stage of the disease, and shorter overall survival. Interestingly, in 2018, a previous report showed that soluble CD163 expression in serum was significantly increased in advanced cutaneous melanoma patients who were responders to the nivolumab immunotherapy ^59^.

In addition to these results, we were able to analyze CD163 protein expression in the tumor tissue, which was correlated with immunotherapy information. This added another layer to our understanding of proteins with predictive values in immunotherapy response.

Furthermore, MTSS2, was found to be significantly downregulated in patients with better therapy response. However, we have not seen a significant correlation between the expression of MTSS2 and survival in immunotherapy cohorts with transcriptomic data. MTSS2, previously named MTSS1L ^60^ is expressed in the central nervous system (CNS) ^61^ and is part of the MTSS protein family ^62^. Despite this, we have limited information about the role of the MTSS2 protein in melanoma progression or therapy response, Hubert et al. suggested that the MTSS2 gene might play a role in cancer susceptibility ^63^.

MTSS2 belongs to the subgroup of I-BAR (Bin/amphiphysin/Rvs) domain-containing proteins ^64^ similar to MTSS1. Notably, MTSS1 has a closer association with melanoma progression where it plays a pivotal role in driving melanocyte metastasis, and elevated MTSS1 expression identifies a subgroup within primary melanomas associated with adverse prognosis ^65^. (**Figure 6**)

**Figure 6.**
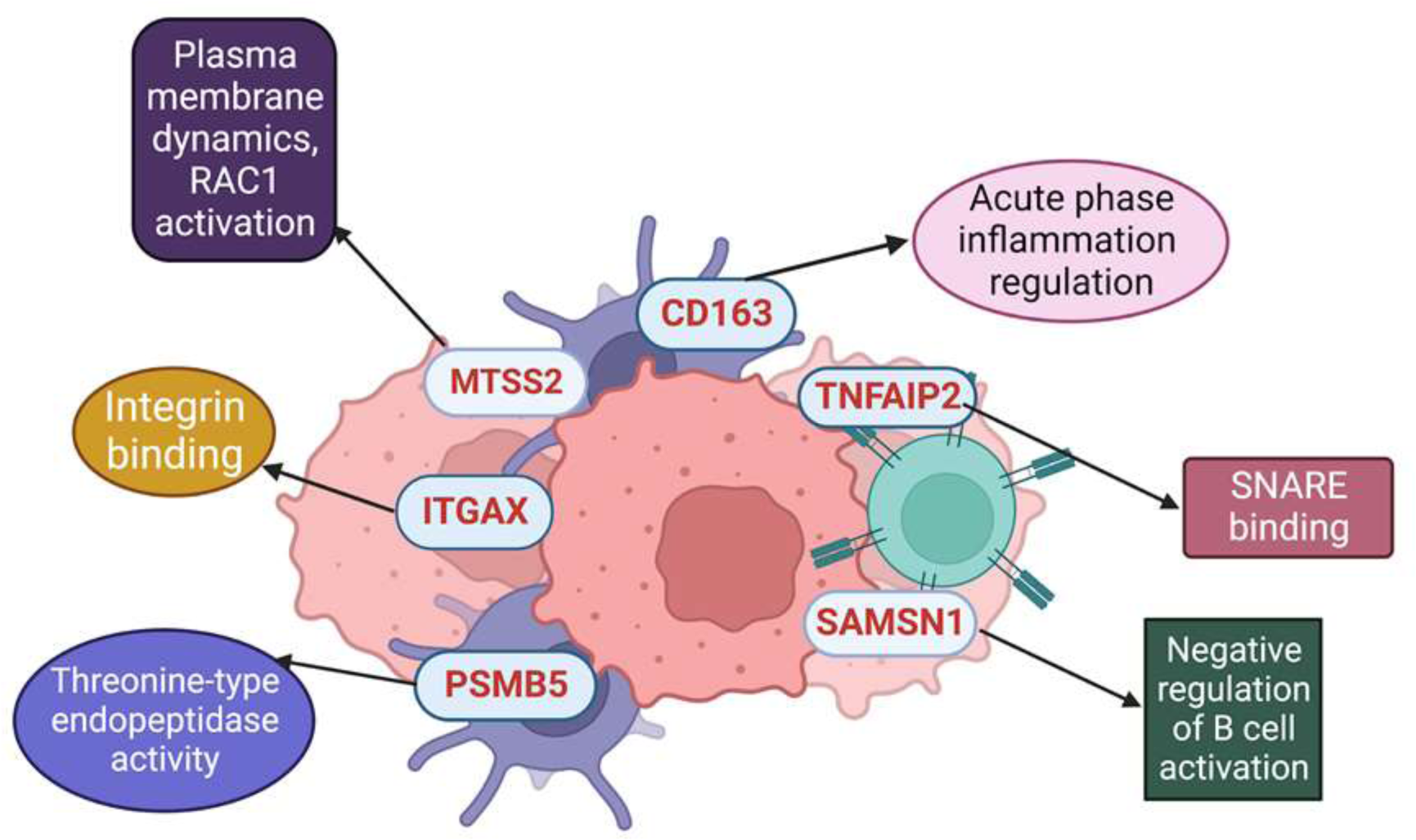
Different molecular mechanisms where the selected proteins are involved.

Moreover, it is noteworthy to mention that among the 6 identified proteins in our findings, two were associated with immune mechanisms (SAMSN1 and CD163), while another two played roles in functions in the tumor microenvironment (TNFAIP2 and ITGAX). The machine-based algorithm (EaSIeR) utilized these pathways to assess the likelihood of responsiveness to immunotherapy.

In conclusion, we highlight for the first time an analysis of one of the largest proteomic datasets in melanoma, searching for predictors which may be associated with immunotherapy response. Through a comprehensive analysis of more than 200 samples from both treated and untreated patients, ranked by a well-defined scoring system, we have identified six candidate proteins.

These six identified proteins as potential biomarkers have been studied across three different metastatic patient cohorts. They exhibit significant correlations with immunotherapy response when evaluated through modelling, as well as independent associations irrespective of other clinical parameters. Moreover, we were able to validate the indicated proteins in various immunotherapy transcriptomic datasets.

The necessity for well-defined biomarkers capable of predicting immunotherapy response as well as survival, and disease progression has reached a critical juncture in the realm of melanoma patient care. Our findings showed functional relationships that some of these biomarkers have with the stroma (*e.g*., ITGAX, PSMB5, TNFAIP2, and MTSS2). Others exhibit stronger connections with immune cells (e.g., CD163, SAMSN1). These proteins hold the promise for sparkling further investigations and may serve as foundation for advancing diagnostics, guiding tailored therapy decisions, aiding in personalized decisions, and ultimately enhancing the life expectancy of metastatic melanoma patients.

## Declare of interests

The authors declare no competing interests.

## Ethical approval statement

### Institutional Review Board Statement

Cohort 1 was conducted according to the guidelines of the Declarations of Helsinki and approved by the Hungarian Ministry of Human Resources, Deputy State Secretary for National Chief Medical Officer, Department of Health Administration. The protocol code is MEL-PROTEO-001, the approval number is 4463- 6/2018/EÜIG and the date of approval is 12 March 2018. Due to the retrospective anonymized FFPE samples, informed consent was not applicable, referring to the MEL-PROTEO-001, 4463-6/2018/EÜIG ethical approval. Cohort 2 was approved by the Regional Ethical Committee at Lund University, Southern Sweden (DNR 191/2007, 101/2013 (BioMEL biobank), 2015/266 and 2015/618). All patients provided written informed consent. The study has been performed in compliance with GDPR. Cohort 3 was carried out in strict accordance with the Declarations of Helsinki and was approved by the Semmelweis University Regional and Institutional Committee of Science and Research Ethics (IRB, SE TUKEB 114/2012 and SE IKEB 191-4/2014). Samples were obtained from the Department of Dermatology and Venereology, Semmelweis University, Budapest, Hungary, under informed consent and a clinical protocol.

## Data and code availability

All original code is available in this paper’s supplemental information (**Supplemental material 3 and Table S3E**) and the scripts used for the statistics are available at https://github.com/indirapla/MM500_ImmuneResponse Github repository ^66^. The data that support the findings of this study are openly available in ProteomeXchange at http://www.proteomexchange.org/ ^67^, reference numbers PXD001725, PXD001724, PXD009630, PXD017968, and PXD026086. The TCGA data was downloaded from cBioPortal https://www.cbioportal.org ^68^. The code for Cohort 2 and 3 can be found at https://github.com/rhong3/TCGA_melanoma ^66^.

## Funding

This study was supported by grants from the Berta Kamprad Foundation, Lund, Sweden. The generous support by Thermo Scientific, Liconic UK, and Tecan was absolutely key for this study. This work was done under the auspices of a Memorandum of Understanding between the European Cancer Moonshot Center in Lund and the U.S. National Cancer Institute’s International Cancer Proteogenome Consortium (ICPC). ICPC encourages international cooperation among institutions and nations in proteogenomic cancer research in which proteogenomic datasets are made available to the public. This work was also done in collaboration with the U.S. National Cancer Institute’s Clinical Proteomic Tumor Analysis Consortium (CPTAC). Further support came from the National Cancer Institute (NCI) CPTAC grants U24CA210972 to D.F. L.V.K. is a recipient of the János Bolyai Research Scholarship of the Hungarian Academy of Sciences and is supported by the Hungarian National Research, Development and Innovation Office (OTKA FK138696) and Semmelweis University STIA-KFI2021 grants. I.N.B. was supported by the Hungarian Academy of Sciences, the grant of OTKA-125509. L.S. was supported by the ÚNKP-21-3-SZTE-102 New National Excellence Program of the Ministry for Innovation and Technology from the source of the National Research, Development and Innovation Fund (University of Szeged, Szeged, Hungary). G.D. was funded by grants (CNPq 440613/ 2016-7; 308241/2019-8) and FAPERJ (E-26/210.173/2 and E-26/201.052/2021) under the Memorandum of Understanding between the Federal University of Rio de Janeiro, Brazil, and Lund University, Sweden. S.K. is acknowledging the support of the Hungarian National Research, Development and Innovation Office (OTKA) support (N-OTKA 114-460).

## Author Contributions

Study Conception & Design, A.S., L.S., A.B.,G.D., L.V.K., I.P.; Performed Experiment or Data Collection, L.H.B, J.G., M.K., J.R.M., A.S., L.S., I.P, A.B., N.W., N. A., J.G., F.N.,V.D., Z. U.,Y.K., Z.P.,T.P.; Computation & Statistical Analysis, L.H.B.,I.P.,B.S.,L.S.,A.B., A.S.,L.V.K, A.L, D.P., A.S.L.; Data Interpretation & Biological Analysis, L.H.B., I.P., L.V.K., A.L., L. S., B.S., A. S., A.B., M.K, J.R.M, Writing – Original Drafts, L.S., A.B., F.N., G.D., A.S., I.P., L.V.K., E.W., A.L., B.G.; Writing – Review & Editing: L.S., A.B., F.N., D.F., G.D., A.S., I.P., L.V.K., A.L., E.W., D.F., K.P., P. H., B.G., G.M-V.; Supervision, G.D., A.S., J.G., I.B.N., L.V.K., E.W., D.F., P.H., S.K., K.P., G.M-V., Administration: Á.J:J, B.B., C. W., A.M.S., J.M., G.M-V.,H.-J. K., R.A. All authors have read and agreed to the published version of the manuscript.

## Limitations

There is no information about the applied treatments of the two untreated cohorts (Cohort 2 and Cohort 3).

## Notes

### Competing Interest Statement

The authors have declared no competing interest.

