## Supplemental Figures for "Predicting immune checkpoint therapy response in three independent metastatic melanoma cohorts"

Figure S1

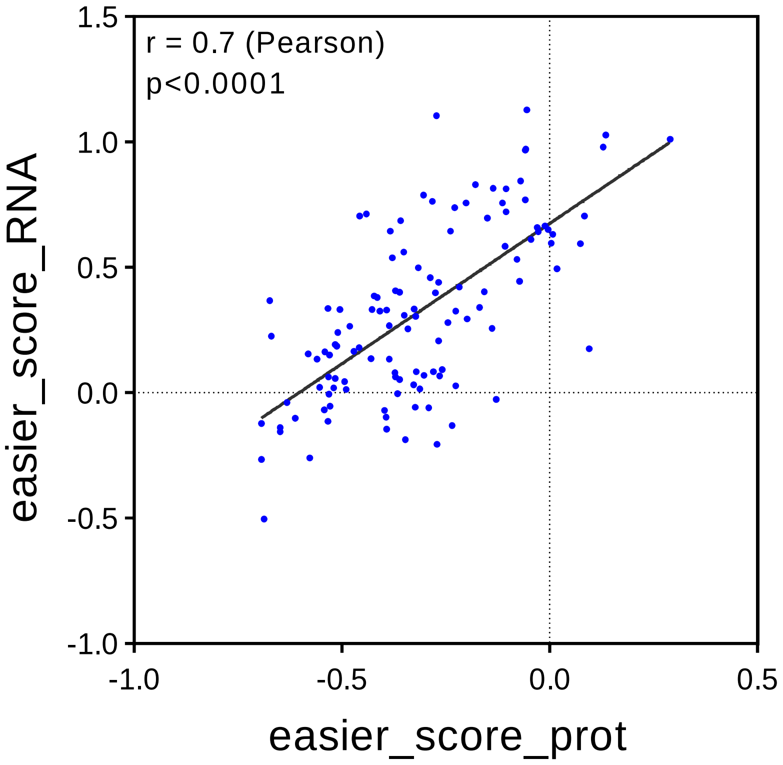

**Figure S1 Comparison of the transcriptomic and proteomic data of EaSIeR score. Related to Figure 2.**

Significant correlation between the EaSIeR score of RNA data and the EaSIeR score of proteomic data supporting the adequate usage of the EaSIeR score algorithm in our study. The input was the protein expression of samples from Cohort 2. The output was the EaSIeR scores for every sample, used for predictors in Cohort 1. Each blue point shows a melanoma sample from Cohort 2.

### Table S1

| Gene | Protein | RNA expression at single cell level in MM cell (nTPM) | Pearson test on ITR score corr. (P-value) | Q1-Q4 Student's T Test (P-value) |
|---|---|---|---|---|
| ITGAX | P20702 | yes (1.8) | **p<0.001 | *p<0.05 |
| TNFAIP2 | Q03169 | yes (30.2) | *p<0.05 | *p<0.05 |
| SAMSN1 | Q9NSI8 | yes (21.4) | *p<0.05 | *p<0.05 |
| CD163 | Q86VB7 | no (0) | *p<0.05 | *p<0.05 |
| PSMB5 | P28074 | yes (115.6) | **p<0.001 | *p<0.05 |
| MTSS2 | Q765P7 | no (0) | *p<0.05 | *p<0.05 |

**Table S1 Summary of the correlations, ITR scores, RNA expression of the identified proteins. Related to Figure 2.**

It represents a summary table about the nTPM values of the identified proteins, the Pearson test correlation on ITR score and the comparison of Q1-Q4 values. (nTMP values mean the RNA expression at single cell level).

### Figure S2

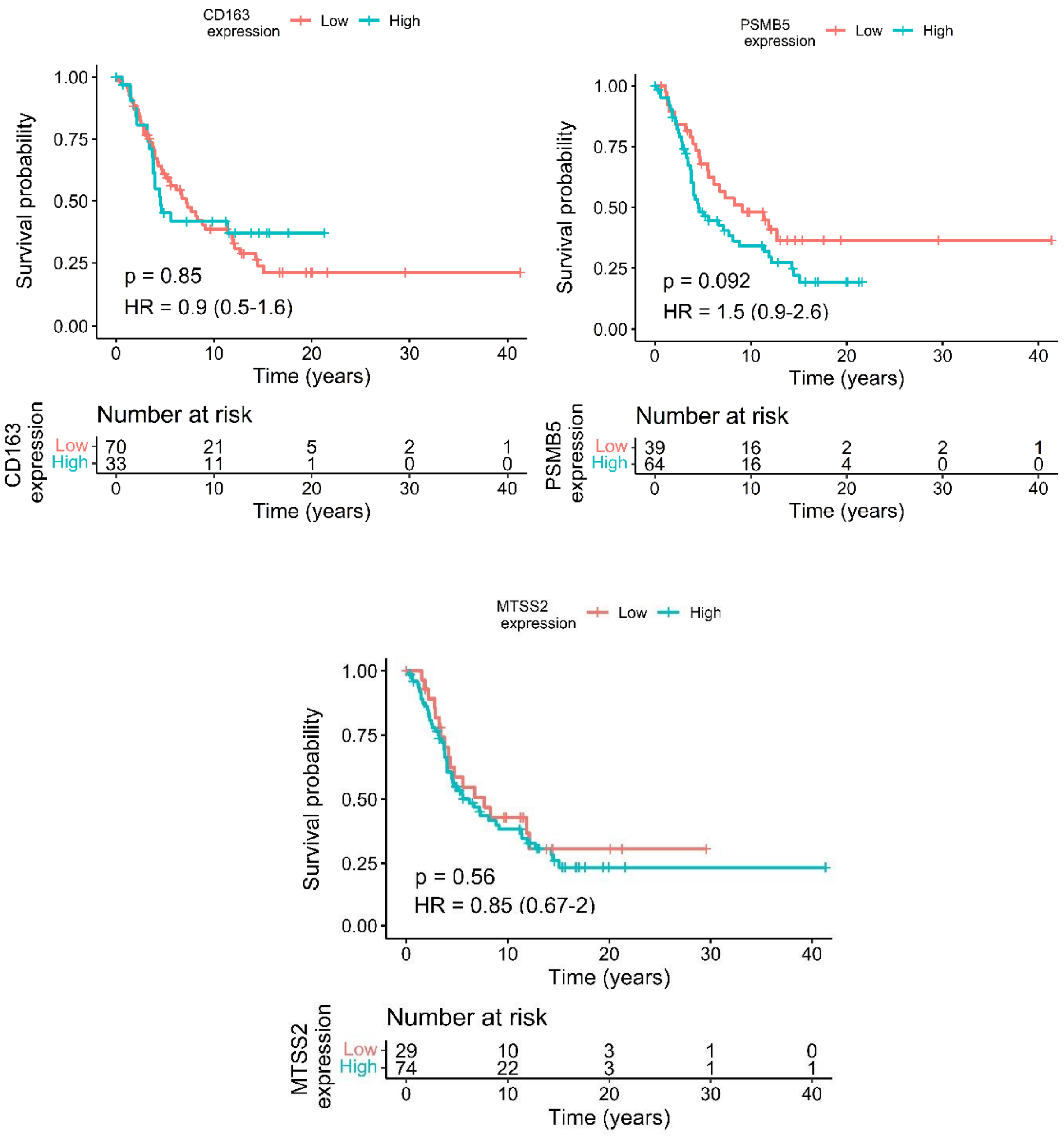

**Figure S2 The survival analysis of PSMB5, MTSS1L and CD163. Related to Figure 3.**

It represents the KM survival plots of PSMB5, MTSS1L and CD163 that was not mentioned in the text.

Figure S3

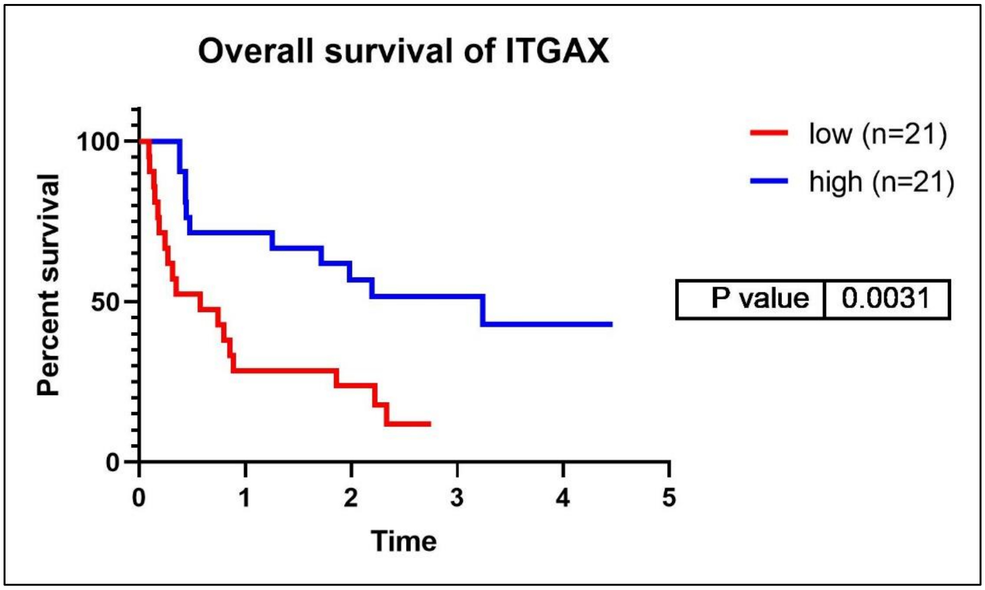

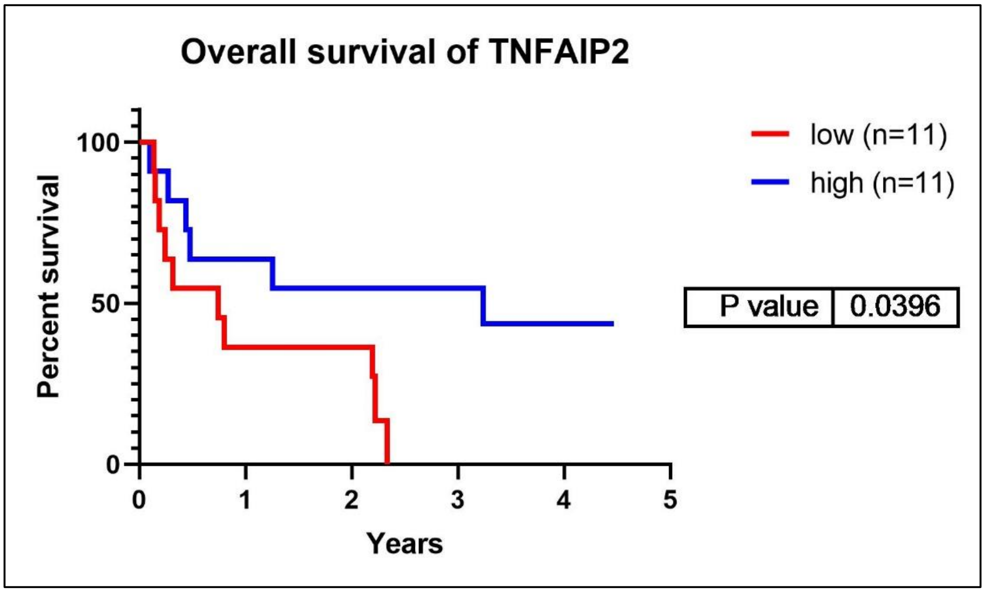

**Figure S3 The overall survival analysis of TNFAPI2 and ITGAX in the transcriptomic validation cohort. Related to Figure 4.**

The figure represents Kaplan-Meier survival analyses of the significantly different expressed genes of the identified proteins for overall survival in the validation CTLA4 immunotherapy transcriptomic cohort. The cut off values are the following: ITGAX – OS:50%; TNFAIP2 – OS:25%; (OS /Overall survival/).
