## Supplemental material 3 for "Predicting immune checkpoint therapy response in three independent metastatic melanoma cohorts": Supplemental Material 3.html

easier score - Segundo data


Code 

- Show All Code
- Hide All Code
- Download Rmd

### easier score - Segundo data

###### Authors: Indira Pla

- Aims
- (1) Data
  cleaning
- (2) Data
  analysis
  - easier score analysis

#### Aims

overall:

To read more about the algorithm behind the ‘easier’ R package you
can follow this link: https://bioconductor.org/packages/release/bioc/vignettes/easier/inst/doc/easier\_user\_manual.html#1\_Introduction
—————————————————————-

**Load libraries:**


```
# packages to load
.packages = c('BiocManager','devtools','tidyverse','AnnotationDbi', 'org.Hs.eg.db','renv','easier')
```


**Load files:**


```
# (1) protein expression data 

prot.exp <- as.data.frame(readxl::read_excel("data/Protein expression_119pat.xlsx")) # Open the data
rownames(prot.exp) <- prot.exp$Accession
```

#### (1) Data cleaning


```
# Filtering by number valid values

perc.VV <- round((30*119)/100,0) # defining the 30% of the samples (considering 119 samples)

prot.exp1 <- subset(prot.exp,prot.exp$`valid values`>perc.VV)
```


**Defining matrix for protein annotations and matrix of protein
expressions**


```
prot.annotation <- prot.exp[,c("Accession","Description","Gene")]
prot.exp_short <- prot.exp1[,5:ncol(prot.exp1)]
```


**converting NA in 0**


```
prot.exp_short1 <- 2^prot.exp_short

prot.exp_short1.imp <- prot.exp_short1
for (i in 1:nrow(prot.exp_short1.imp)) {
  for (j in 1:ncol(prot.exp_short1.imp)) {
    ifelse(is.na(prot.exp_short1.imp[i,j]),
           prot.exp_short1.imp[i,j]<-0,
           prot.exp_short1.imp[i,j]<-prot.exp_short1.imp[i,j])
  }
}
```


**Checking and removing duplicated genes**


```
prot.exp_short1.imp$Accession <- rownames(prot.exp_short1.imp)

prot.exp_short.imp1 <- plyr::join_all(list(prot.exp_short1.imp,prot.annotation), by = 'Accession')
```


If there is no Gene symbol, it will be substituted by the accession
of the protein


```
for (i in 1:nrow(prot.exp_short.imp1)) {
  if(is.na(prot.exp_short.imp1$Gene[i]))prot.exp_short.imp1$Gene[i]<- prot.exp_short.imp1$Accession[i]
}

dup <- prot.exp_short.imp1[duplicated(prot.exp_short.imp1$Gene),c('Accession','Gene')]

dup1<-prot.exp_short.imp1[prot.exp_short.imp1$Gene %in% dup$Gene,c('Accession','Gene')]

# removing duplicated genes
rownames(prot.exp_short.imp1a) <- prot.exp_short.imp1a$Gene
```


**Final curated protein expression data**


```
# removing "Accession","Description","Gene" columns 
prot.exp_short.imp1b <- prot.exp_short.imp1a %>% dplyr::select(-c("Accession","Description","Gene" ))
```

#### (2) Data analysis

##### easier score analysis

###### 2.1 Selecting cancer type


```
# Cancer type 
cancer_type_MM <- 'SKCM'   # skin cutaneous melanoma (SKCM)
```

###### 2.2 Compute hallmarks of immune response


```
library(easier)
hallmarks_of_immune_response <- c("CYT", "Roh_IS", "chemokines", "Davoli_IS", 
                                  "IFNy", "Ayers_expIS", "Tcell_inflamed",
                                  "RIR", "TLS")


immune_response_scores_MM <- compute_scores_immune_response(RNA_tpm = prot.exp_short.imp1b, selected_scores = hallmarks_of_immune_response)
```


```
head(immune_response_scores_MM)
```


```
# Saving table in the computer 
# write.csv2(immune_response_scores_MM,'output/Proteom_2do_immune_response_scores_MM.csv')
```

###### 2.3 Compute quantitative descriptors of the TME

**Cell Fractions**


```
cell_fractions_MM <- compute_cell_fractions(RNA_tpm = prot.exp_short.imp1b)
```


```
Running quanTIseq deconvolution module

Gene expression normalization and re-annotation (arrays: FALSE)

Removing 17 noisy genes

Removing 15 genes with high expression in tumors

Signature genes found in data set: 88/138 (63.77%)

Mixture deconvolution (method: lsei)

Deconvolution successful!
Cell fractions computed!
```


```
head(cell_fractions_MM)
```


```
# Saving table in the computer
#write.csv2(cell_fractions_MM,'output/Proteom_2do_cell_fractions_MM.csv')
```


**Pathway activity**

Applying PROGENy (Holland, Szalai, and Saez-Rodriguez 2020; Schubert
et al. 2018) method to count data from RNA-seq, the activity of 14
signaling pathways

BUT WE DONT HAVE RNA COUNT MATRIX\*\*\*\*\*\*\* WE COULD GET IT FROM THE
TRANSCRIPTOMIC GROUP


```
#-----Pathway activity------#
# pathway_activities_MM <- compute_pathway_activity(RNA_counts = RNA_counts,
#                                                remove_sig_genes_immune_response = FALSE,
#                                                verbose = TRUE)
# head(pathway_activities_MM)
```


By applying DoRothEA (Garcia-Alonso et al. 2019) method to TPM data
from RNA-seq, the activity of 118 transcription factor (TF) can be
inferred as follows:


```
tf_activities_MM <- compute_TF_activity(RNA_tpm = prot.exp_short.imp1b)
```


```
Regulated transcripts found in data set: 2161/3254 (66.4%)
Registered S3 method overwritten by 'data.table':
  method           from
  print.data.table     
TF activity computed!
```


```
head(tf_activities_MM[,1:5])
```


```
#write.csv2(tf_activities_MM,'output/Proteom_2do_TFs_activities_MM.csv')
```


**Ligand-receptor (LR)**


```
#----- Ligand-receptor (LR)-----#

lrpair_weights_MM <- compute_LR_pairs(RNA_tpm = prot.exp_short.imp1b,
                                   cancer_type = "pancan")
```


```
LR signature genes found in data set: 455/644 (70.7%)
Ligand-Receptor pair weights computed
```


```
head(lrpair_weights_MM[,1:5])
```


```
#write.csv2(lrpair_weights_MM,'output/Proteom_2do_lrpair_weights_MM.csv')
```


Using the ligand-receptor weights as input, 169 cell-cell interaction
scores can be derived as in the chunk below.


```
ccpair_scores_MM <- compute_CC_pairs(lrpairs = lrpair_weights_MM, 
                                  cancer_type = "pancan")
```


```
CC pairs computed
```


```
# CC pairs computed
head(ccpair_scores_MM[,1:5])
```


```
#write.csv2(ccpair_scores_MM,'output/Proteom_2do_ccpair_scores_MM.csv')
```

###### 2.4 Obtain patients’ predictions of immune response


```
predictions_MM <- predict_immune_response(immunecells = cell_fractions_MM,
                                          #pathways = pathway_activities_MM,
                                          tfs = tf_activities_MM,
                                          lrpairs = lrpair_weights_MM,
                                          ccpairs = ccpair_scores_MM,
                                          cancer_type = cancer_type_MM, 
                                          verbose = TRUE)
```


Once we obtained patients’ predicted immune response, two different
scenarios should be considered in which:

*-patient\_response is known and therefore the accuracy of easier
predictions can be evaluated* *-patient\_response is unknown and
no assessments can be carried out*

What if I have an immunotherapy dataset where patients’ response is
not available? In this likely scenario, an score of likelihood of immune
response can be assigned to each patient by omitting the argument
patient\_response within the function assess\_immune\_response.


```
output_eval_no_resp_MM <- assess_immune_response(predictions_immune_response = predictions_MM,
                                                 RNA_tpm = prot.exp_short.imp1b,
                                                 # TMB_values = TMB,
                                                 weight_penalty = 0.5)
```


```
Scoring patients' as real response is not provided

Error in aggregate.data.frame(lhs, mf[-1L], FUN = FUN, ...) : 
  no rows to aggregate
```


**Figure 1** output: Boxplot of patients’ easier score
showing its distribution across the 10 different tasks. **Figure
2** output: Scatterplot of patients’ prediction when combining
easier score with tumor mutational burden showing its distribution
across the 10 different tasks.


```
output_eval_no_resp_MM[[1]]
output_eval_no_resp_MM[[2]]
```

###### 2.5 Retrieve easier scores of immune response


```
easier_derived_scores_MM <- retrieve_easier_score(predictions_immune_response = predictions_MM,
                                                  # TMB_values = TMB,
                                                  easier_with_TMB = c("weighted_average", 
                                                                      "penalized_score"),
                                                  weight_penalty = 0.5)

head(easier_derived_scores_MM)
```


```
#write.csv(easier_derived_scores_MM,'output/easier_derived_scores_prot_119MM.csv')

hist(easier_derived_scores_MM$easier_score, breaks = 50)
```


```
shapiro.test(easier_derived_scores_MM$easier_score)
```


```
    Shapiro-Wilk normality test

data:  easier_derived_scores_MM$easier_score
W = 0.98234, p-value = 0.1206
```

###### 2.6 Interpret response to immunotherapy through systems biomarkers


```
output_biomarkers <- explore_biomarkers(immunecells = cell_fractions_MM,
                                        #pathways = pathway_activities_MM,
                                        tfs = tf_activities_MM,
                                        ccpairs = ccpair_scores_MM,
```


```
Warning: Removed 3 rows containing non-finite values (`stat_boxplot()`).Warning: Removed 3 rows containing missing values (`geom_point()`).
```
